# Oligodendrocyte Development and Myelination in the 22q11.2 Deletion Syndrome

**DOI:** 10.64898/2026.11.17.767424

**Authors:** Amelia Hutchin, Mariyam Lokhand, Caitlyn Rock, Allison Yee, Megan Baldemor, Regina Lara, Zane Turner, Sahej Sidhu, Laura Cocas

## Abstract

The 22q11.2 deletion syndrome is associated with a high risk of neurodevelopmental disorders, including a 25% chance of developing schizophrenia (SCZ). SCZ is prevalent in 0.4-0.7% of the population. Evidence suggests that the cognitive deficits of SCZ may be caused by neurodevelopmental disconnection due to impaired oligodendrocyte (OL) maturation and myelination. We hypothesized that the 22q11.2 deletion syndrome impairs OL life cycle progression, halting oligodendrocytes in their immature phase and preventing advancement to the mature stage. This in turn creates myelination changes, demonstrated by a decrease in myelin. To test this, we utilized a 22q11.2 deletion mouse model focusing on the prefrontal cortex (PFC), a late-myelinating brain region. We evaluated PFC regions (mPFC, dlPFC, mOFC, lOFC) in adolescent (P14 and P28) mutant mice and controls. Additionally, we examined the two-hit hypothesis of schizophrenia using a maternal separation model. At P28, mutant mice displayed a significant increase in immature OLs and mature OLs in the PFC, and a decrease in myelination and myelin process length compared to controls. At P14, mutant mice showed subtle reductions in oligodendrocyte precursor cells and subtle myelination alterations in the mPFC. When considering the two-hit hypothesis, we saw equivalent reductions in myelination in the genetic-only, environmental-only, and two-hit groups. Together, these data demonstrate that the 22q11.2 deletion impairs OL maturation and functional myelination between P14 to P28, a critical window for neurodevelopment. Examining the impact of SCZ on OLs could reveal key intervention targets before SCZ symptom onset, and may determine the role of OLs and the 22q11.2 deletion in neural disconnectivity in schizophrenia.

## Introduction

Schizophrenia is a neurodevelopmental psychiatric disorder characterized by positive (hallucinations and delusions), negative (avolition and anhedonia), and cognitive symptoms (decreased working memory and executive dysfunction). There are many well-documented genetic risk factors for schizophrenia, including DiGeorge’s syndrome^1^. 22q11.2 deletion syndrome (22q11.2DS), also known as DiGeorge’s syndrome, is a genetic disorder caused by a hemizygous deletion on chromosome 22^2^. Previous studies have identified widespread white matter abnormalities in individuals with 22q11.2DS^3^. 22q11.2DS occurs in approximately 1 in 2,000–4,000 live births^4^, while approximately 25% of individuals with 22q11.2DS develop schizophrenia^5^. There is experimental evidence that reduced expression of *Tbx1*, a transcription factor gene in the deleted region of 22q11.2DS, affects the composition of myelinated axons^6^. Additionally, single-cell RNA sequencing of the prefrontal cortex in 22q11.2DS mice revealed transcriptional alterations in oligodendrocyte-lineage cells^7^. However, the effects of this deletion on oligodendrocyte-lineage development and myelination during adolescence are not well understood. Adolescence represents a critical developmental period in 22q11.2DS, with one longitudinal study reporting that 82.6% of individuals who developed psychotic disorders experienced symptom onset between 14 and 22 years of age^8^. Understanding the effects of 22q11.2DS on oligodendrocyte development during adolescence is therefore particularly important, as myelination continues throughout this period, especially in late-maturing regions such as the prefrontal cortex.

Oligodendrocytes are glial cells responsible for producing myelin in the central nervous system, forming sheaths around CNS axons, which enables efficient saltatory conduction of action potentials^9^. Myelination continues throughout adolescence and into early adulthood, particularly in late-maturing association cortical regions such as the PFC^10^,^11^. Oligodendrocyte progenitor cells (OPCs) proliferate to produce cells that progress through intermediate premyelinating stages, and eventually become mature, myelinating oligodendrocytes. This differentiation process is tightly regulated by transcription factors and intracellular signaling pathways that coordinate oligodendrocyte maturation and myelin formation.

Previous studies have demonstrated that disrupting regulatory pathways involved in oligodendrocyte differentiation, including mechanistic target of rapamycin (mTOR) signaling, can impair oligodendrocyte maturation and developmental myelination^12^. Although abnormalities in white matter microstructure have been documented in individuals with 22q11.2DS, there remains an incomplete understanding of how the deletion affects specific stages of oligodendrocyte-lineage development and myelination during adolescence. Experimental evidence suggests that reduced dosage of genes within the 22q11.2 deletion region may contribute to these abnormalities by disrupting oligodendrocyte development and myelin formation^13^. For example, mice heterozygous for Tbx1, a gene located within the 22q11.2 deletion region, exhibited reduced expression of the OPC-associated marker Ng2, decreased oligodendrocyte production from postnatal progenitor cells, and alterations in the composition of myelinated axons in the fimbria^14^.

Schizophrenia is hypothesized to arise from interactions between genetic susceptibility and environmental risk factors, consistent with the two-hit model of neurodevelopmental disorders^15^. This model proposes that genetic vulnerability may alter the brain’s response to subsequent environmental stressors, potentially increasing susceptibility to psychiatric disorders. Maternal separation is an established rodent model of early-life stress that has been shown to disrupt oligodendrocyte development and myelination, particularly in the medial prefrontal cortex (mPFC). Previous work demonstrated that neonatal maternal separation impairs prefrontal myelination and cognitive function in rats through mechanisms involving Wnt signaling^16^. Similarly, maternal separation induces premature oligodendrocyte differentiation in the developing mouse mPFC, followed by depletion of the OPC population in adulthood^17^. Together, these findings suggest that early-life stress alters oligodendrocyte maturation and myelination, modifying the neurodevelopmental consequences of genetic vulnerability associated with 22q11.2DS.

In this study, we sought to investigate the effects of 22q11.2 deletion and early-life stress on oligodendrocyte development and myelination during adolescence. Using Df(h22q11)/+ mice, a genetic model of 22q11.2DS, we examined oligodendrocyte-lineage populations and myelination across postnatal developmental stages P14 and P28. We employed a maternal-separation paradigm to investigate whether early-life stress modifies the effects of genetic vulnerability on oligodendrocyte development. We hypothesized that 22q11.2 deletion and maternal separation would alter oligodendrocyte maturation and myelination, with greater cellular phenotypes in mice exposed to both genetic and environmental risk factors.

## Methods

### Animals

All animal procedures were approved by the Institutional Animal Care and Use Committees at Santa Clara University. We used the Df(h22q11)/+ deletion mouse model (C57BL/6-Del(16Dgcr2-Hira)1Tac; Taconic Biosciences) which has a genomic deletion on chromosome 16 that corresponds to the 22q11.2 region on the human chromosome. The 22q11 deletion in humans indicates high risk for neurodevelopment disorders, including schizophrenia, which was our reasoning for choosing this mouse model. We bred male mice heterozygous for this deletion with wildtype females and genotyped pups using PCR. We evaluated PFC regions (mPFC, dlPFC, mOFC, lOFC) in adolescent (P14 and P28) mutant mice and age-matched wildtype controls, sex-balanced.

### Tissue Processing and Sectioning

Mice were perfused with 4% PFA, brains were harvested and postfixed for 2 hours, and cryoprotected in 30% sucrose overnight. Brains were embedded in O.C.T. compound (Sakura Finetek; Torrance, CA, USA) and stored at ™20 °C, sectioned into 30 µm coronal slices, mounted and stored in PBS with azide at 4 °C until staining.

### Immunohistochemistry

Sections were washed thrice with 0.2 % PBS for 10 min at a time, then incubated in blocking buffer (10 % lamb serum, 0.2 % Triton, 0.005 % sodium azide, and 1:50000 DAPI in PBS) for 30 min, after which they were incubated in primary antibody for 48 hours at room temperature. Following three washes with PBS, sections were then incubated in secondary antibody for 1 hour at room temperature, followed by three washes in PBS, and mounted using Fluoromount-G mounting medium (Thermo Fisher Scientific; Waltham, MA, USA). Stained sections were imaged using a Leica TCS SP5 Confocal Microscope. The following primary antibodies were used: 1:100 anti-Olig2 Rb (Abcam, ab109186); 1:100 anti-Olig2 Ms (Millipore, MABN50), 1:100 anti-MBP rat (Millipore, MAB386), 1:50 anti-CC1 mouse (Calbiochem, OP80), 1:100 anti-rabbit Bcas (Thermo Fisher, PA5-20904), Parvalbumin (Millipore, MAB 1572) and WFA biotinylated (Vector Labs, B-1355-2). All secondary antibodies were Alexa Fluor secondary antibodies from Jackson Immunoresearch, raised in goat, and conjugated to 488, 546, or 633 fluorophores as needed except for Streptavidin (Biotium, CF-29035) secondaries used for WFA antibodies.

### EdU Injections and Staining

We injected animals with EdU 3 hours before sacrificing, using Click-it chemistry to mark EdU-labeled cells and staining for Olig2 to mark the OL-lineage. Sections were stained with EdU using the Click-iT® EdU Imaging Kit with Alexa Fluor® 647 Azide (ThermoFisher, C10337, Waltham, MA, USA) according to manufacturer instructions. Following EdU staining, slides were incubated in blocking buffer (10 % lamb serum, 0.33 % Triton, 0.005 % sodium azide, and 1:50000 DAPI in PBS) for 30 min, then immunostained overnight with 1:50 anti-rabbit Olig2. Slides were washed in PBS three times and mounted using Fluoromount-G mounting medium.

### 24 Hour Maternal Separation

P9 pups were maintained for 24 continuous hours without the dam in a clean cage on a heating pad, and were monitored at separation, 1 hour after setup, every 4 hours during workday hours, and again at reunion. We recorded litter cohesion/huddling, skin color, movement/activity on stimulation, visible dehydration, and deaths, if any. We returned the pups to the dam on P10 and observed the cage continuously for 15–30 minutes to confirm retrieval/nursing/acceptance, and rechecked at 2 hours and 24 hours to look for maternal aggression, cannibalism, failure to nurse, and cold or injured pups. Control litters remained with the dam under standard rearing conditions.

### Data Collection and Statistical Analysis

Sections were analyzed to manually quantify the number of co-labeled cells by category. For all analyses 3 technical replicates were quantified per biological replicate for each brain region quantified; no images with < 2 technical replicates were used. The Leica TCS SP5 Confocal Microscope was used to collect all images. All positive cell labeling was confirmed by a DAPI counterstain. The region of interest was defined for each brain region measured and maintained across all samples. The bregma level was also kept uniform across all samples, with the region of PFC taken from sections between Bregma 2.71 mm and Bregma 1.98 mm. All schematics were created with Affinity.

MBP % area was measured using FIJI to threshold images. MBP immunodensity compared the mean gray value of the background to the mean gray value of the whole image. MBP process lengths were measured using the average of 30 lengths from each technical replica.

We utilized a mixed-model statistical analysis. We first tested for normality using the Shapiro-Wilk test. We analyzed non-parametric data using the Kolmogorov-Smirnov test; otherwise unpaired student’s t-tests were used. Maternal separation data for P28 MBP was analyzed using ANOVA and post-hoc Tukey HSD tests. Descriptive statistics are included in all figure legends. All data was analyzed and plotted using GraphPad Prism version 11.0.02.

## Results

### Effects on OPC proliferation and differentiation

We determined the impact of the 22q11.2 deletion on OPC proliferation and maturation in the adolescent brain at P28. We found no significant difference of EdU+Olig2+ cells in any region of the PFC.(Figure 1).

**Figure 1.**
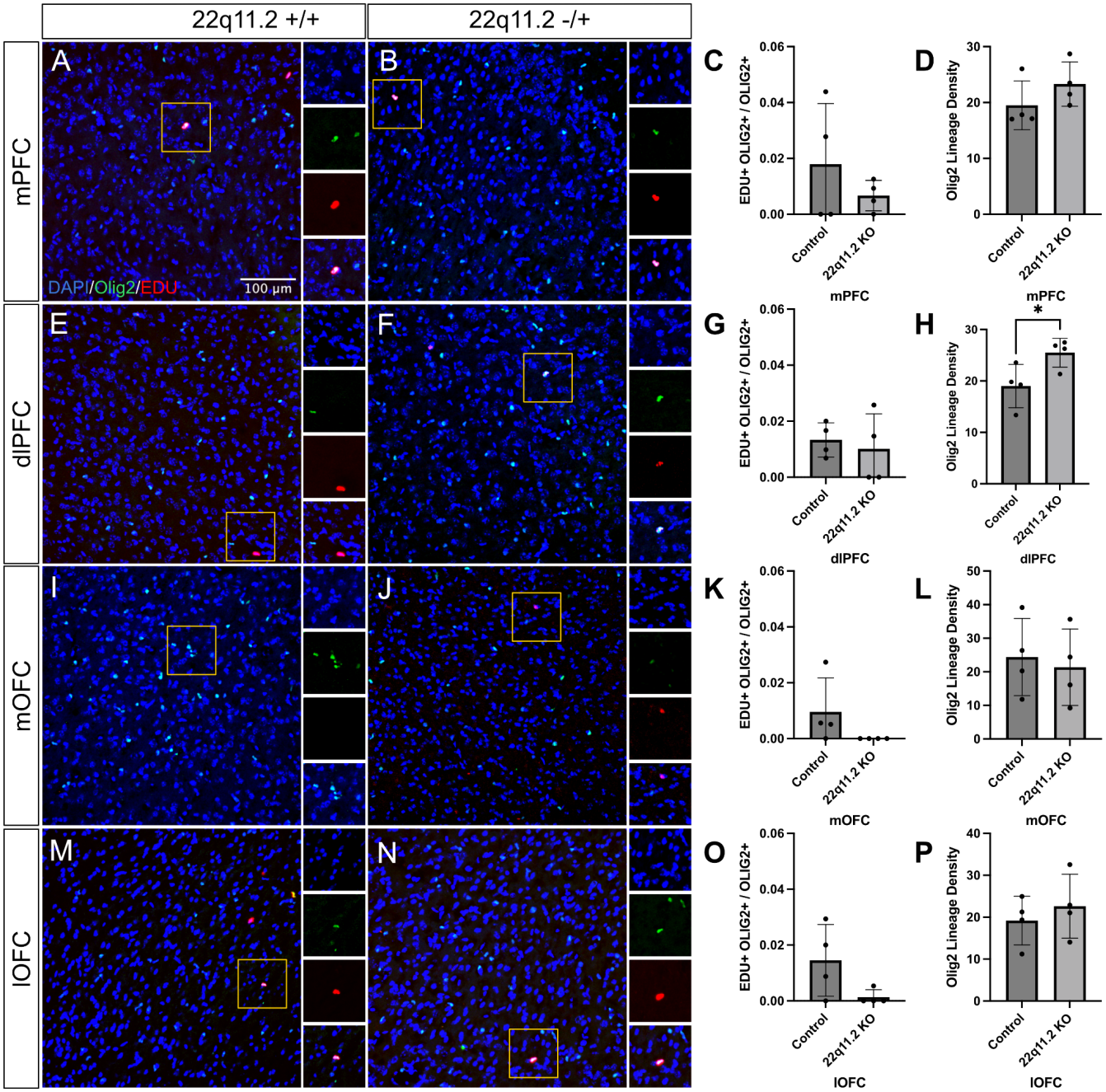
Adolescent 22q11.2 Deletion Mice Have No Change in OPC Proliferation in the Prefrontal Cortex. Representative coronal sections showing DAPI (blue), Olig2 (green), and click-it chemistry EdU (red) staining in P28 WT (n=4) and 22q11.2 KO (n=4) mice across the medial prefrontal cortex (A, B), dorsolateral prefrontal cortex (E, F), medial orbitofrontal cortex (I, J), and lateral orbitofrontal cortex (M, N). (C, G, K, O) Ratio of EdU+Olig2+ cells to total Olig2+ cells. (D, H, L, P) Olig2+ lineage density. Data presented as mean ± SEM, unpaired two-tailed t-test: (C) t=1.006, df=6, p=0.3534; (D) Kolmogorov-Smirnov test d=0.7500, p=0.2286; (G) t=0.4608, df=6, p=0.6612; (H) t=2.561, df=6, p=0.0429; (K) Kolmogorov-Smirnov test d=0.7500, p=0.1429; (L) t=3771, df=6, p=0.7191; (O) Kolmogorov-Smirnov test d=0.7500, p=0.1429; (P) t=0.7167, df=6, p=0.5005.

We then examined the impact of the deletion on the maturation of OLs, looking at the density of immature and mature cells, marked by BCAS1+ and CC1+ cells, respectively. We used Olig2 to mark the OL-lineage. We saw an increase in BCAS1+/Olig2+ cells in the mPFC, dlPFC, and lOFC (Figure 2). This demonstrates an increase in the density of immature cells in several regions of the prefrontal cortex. We did not see a significant increase of immature cells in the mOFC. We next looked at mature oligodendrocytes, CC1+/Olig2+ cells, and saw an increase of mature oligodendrocyte density only in the mPFC (Figure 3).

**Figure 2.**
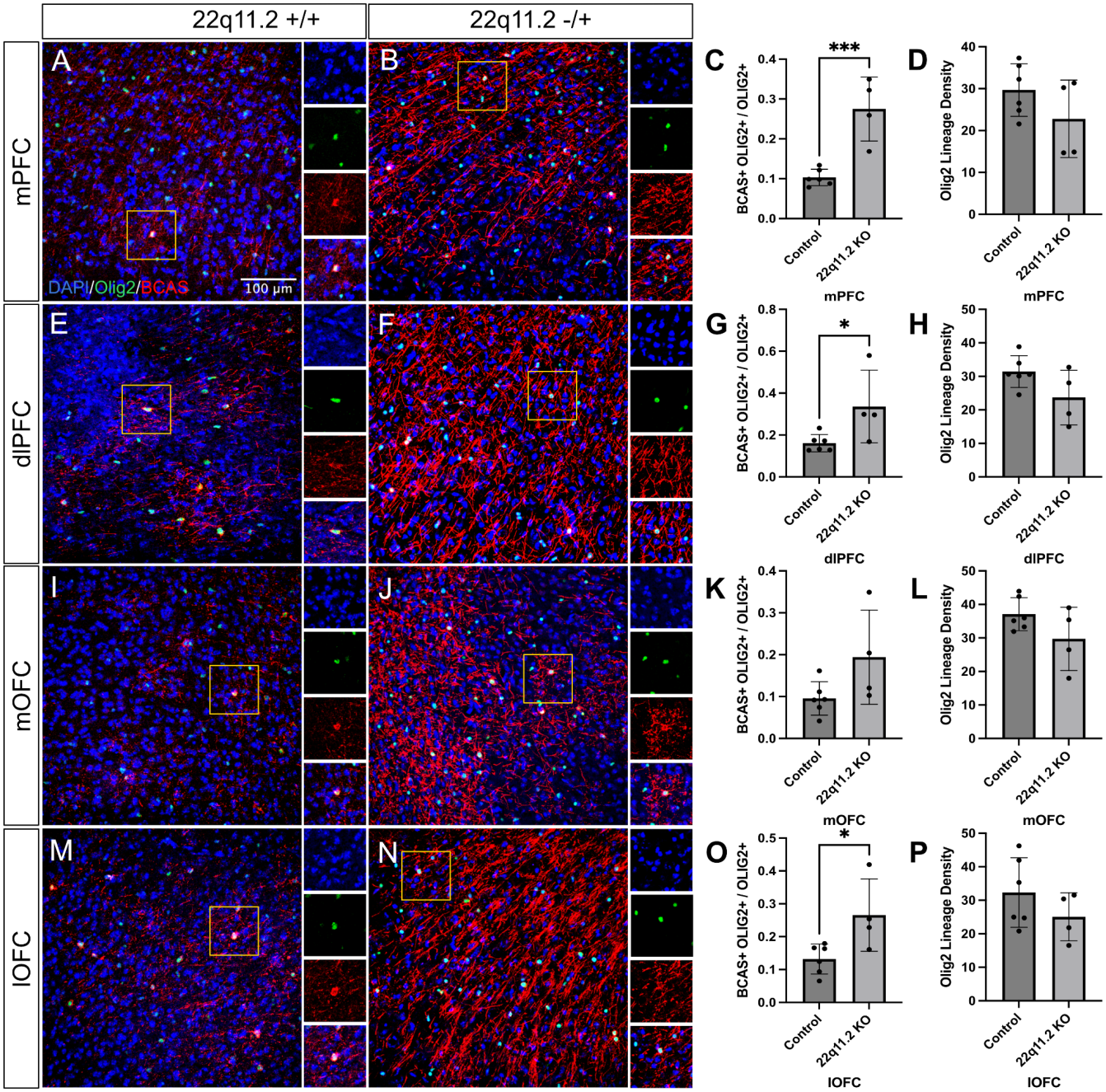
Adolescent 22q11.2 Deletion Mice Have Increased Immature Oligodendrocytes in the Prefrontal Cortex. Representative coronal sections showing DAPI (blue), Olig2 (green), and BCAS (red) staining in P28 WT (n=5) and 22q11.2 KO (n=4) mice across the medial prefrontal cortex (A, B), dorsolateral prefrontal cortex (E, F), medial orbitofrontal cortex (I, J), and lateral orbitofrontal cortex (M, N). (C, G, K, O) Ratio of BCAS+Olig2+ cells to total Olig2+ cells. (D, H, L, P) Olig2+ lineage density. Data presented as mean ± SEM, unpaired two-tailed t-test: (C) t=5.135, df=8, p=0.0009; (D) t=1.414, df=8, p=0.1952; (G) t=2.430, df=8, p=0.0.412; (H) t=1.927, df=8, p=0.0.0902; (K) t=2.014, df=7, p=0.0788; (L) t=1.646, df=8, p=0.1383; (O) t=2.713, df=8, p=0.0265; (P) t=1.210, df=8, p=0.2610.

**Figure 3.**
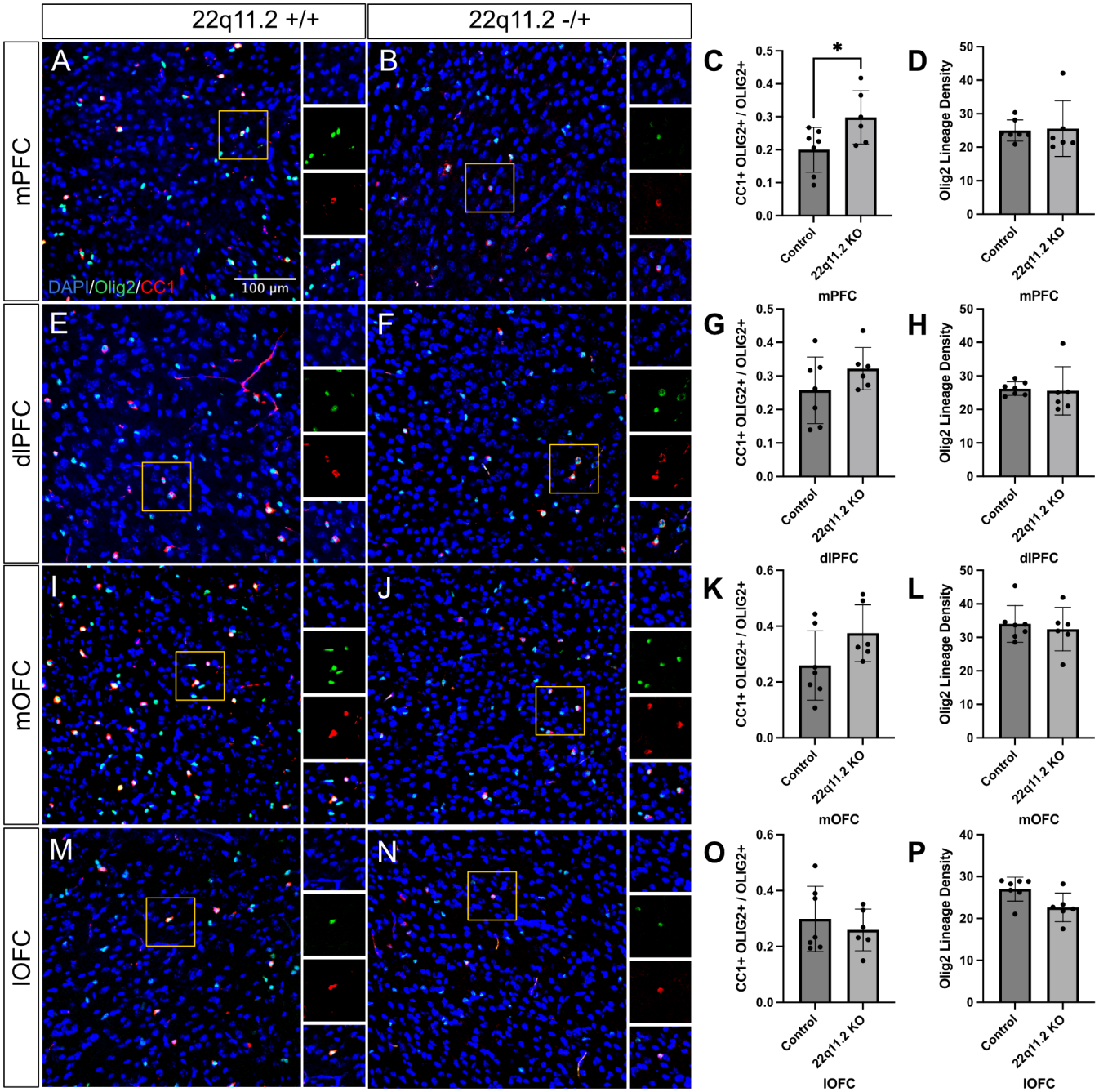
Adolescent 22q11.2 Deletion Mice Have Increased Mature Oligodendrocytes in the Medial Prefrontal Cortex. Representative coronal sections showing DAPI (blue), Olig2 (green), and CC1 (red) staining in P28 WT (n=7) and 22q11.2 KO (n=6) mice across the medial prefrontal cortex (A, B), dorsolateral prefrontal cortex (E, F), medial orbitofrontal cortex (I, J), and lateral orbitofrontal cortex (M, N). (C, G, K, O) Ratio of CC1+Olig2+ cells to total Olig2+ cells. (D, H, L, P) Olig2+ lineage density. Data presented as mean ± SEM, unpaired two-tailed t-test: (C) t=2.288, df=11, p=0.0360; (D) Kolmogorov-Smirnov test d=0.5238, p=0.2774; (G) t=1.370, df=11, p=0.1979; (H) Kolmogorov-Smirnov test d=0.5000, p=0.3328; (K) t=1.815, df=11, p=0.0.969; (L) t=0.4762, df=11, p=0.6431; (O) t=0.7083, df=11, p=0.0.4935; (P) Kolmogorov-Smirnov test d=0.6905, p=0.0676.

### Effects on myelination

Since we found an increase in immature and mature OLs, we hypothesized that we would find a concordant increase in myelination in the PFC. However, in all regions of the PFC we observed a significant decrease in myelination, as measured by both area fraction and immunodensity of MBP labeled myelin. Additionally, we measured myelin process lengths, and found significantly decreased lengths within the brains of mice with the 22q11 deletion (Figure 4). This suggests that there is a mechanism leading to decreased myelination that is unrelated to OL cell density.

**Figure 4.**
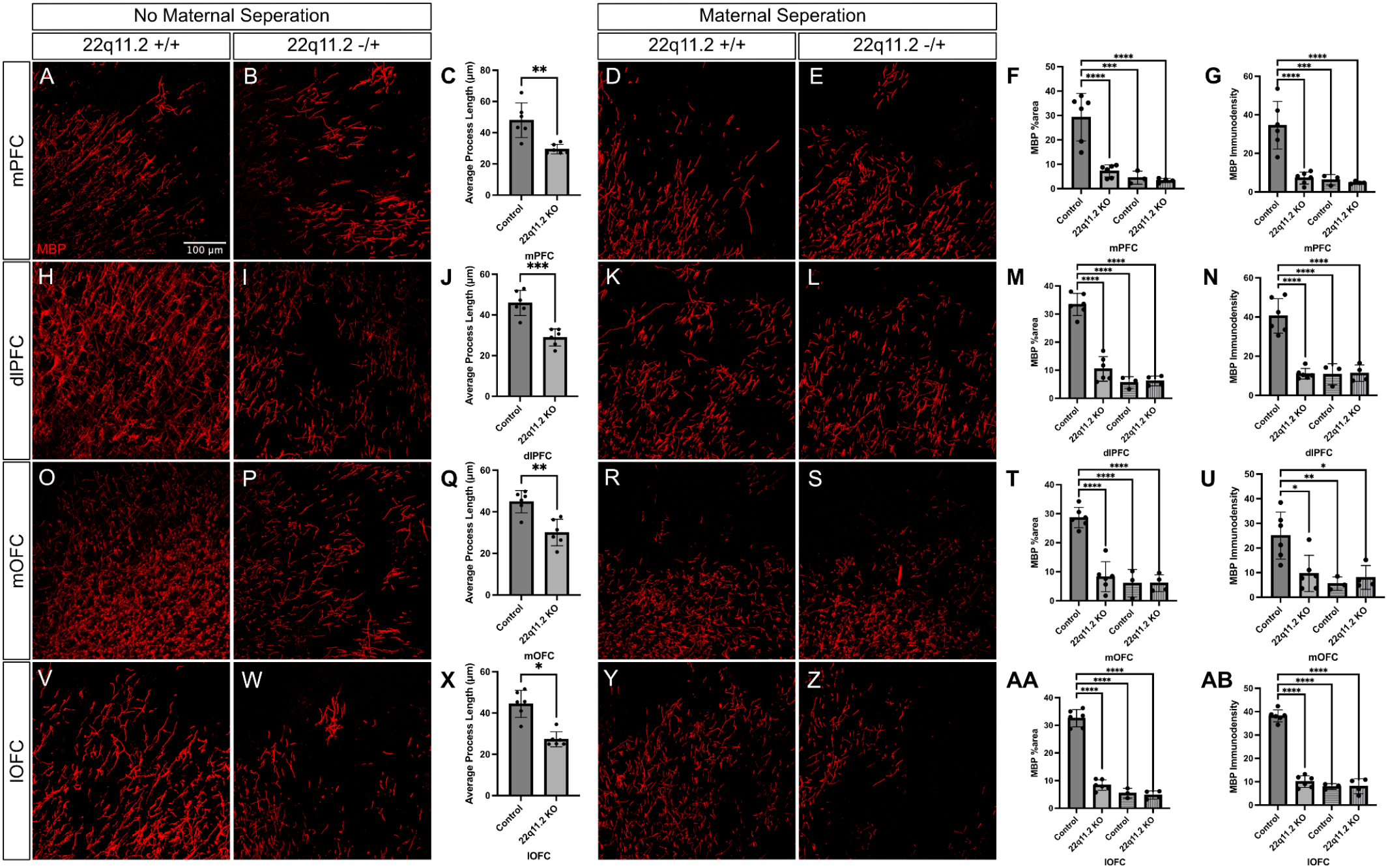
22q11.2 Deletion and Maternal Separation Two-Hit Model Show Reduction in Myelination in the Adolescent Prefrontal Cortex. Representative coronal sections showing MBP (red) staining in P28 WT (n=6), 22q11.2 KO (n=6), maternally separated WT (n=3), and maternally separated KO (n=4) mice across the medial prefrontal cortex (A, B, D, E), dorsolateral prefrontal cortex (H, I, K, L), medial orbitofrontal cortex (O, P, R, S), and lateral orbitofrontal cortex (V, W, Y, Z). (C, J, Q, X) MBP average process length (μm) for non-separated mice, unpaired two-tailed t-test: (C) t=3.926, df=10, p=0.0028; (J) t=5.563, df=10, p=0.0002; (Q) t=4.343, df=10, p=0.0015; (X) Kolmogorov-Smirnov test d=0.8333, p=0.0260. (F, M, T, AA) MBP area fraction. (G, N, U, AB) MBP immunodensity. One-way ANOVA with Tukey’s HSD for pairwise comparisons: (F) F(3, 15)=22.99, p<0.0001; (G) F(3, 15)=20.08, p<0.0001; (M) F(3, 15)=70.81, p<0.0001; (N) F(3, 15)=33.39, p<0.0001; (T) F(3, 15)=36.47, p<0.0001; (U) F(3, 15)=7.379, p=0.0029; (AA) F(3, 15)=117.0, p<0.0001; (AB) F(3,15)=179.5, p<0.0001. Maternally separated groups marked by stripped columns. Data presented as mean ± SEM. * p<0.05, ** p<0.01, *** p<0.001, **** p<0.0001

### Effects from a two-hit model

We next examined the impact of genetics and environmental stressors on OL myelination. We used a maternal separation paradigm, which involved a 24hr separation of pups from their mother at P9. Surprisingly, we found a reduction in myelination after maternal separation alone, compared to controls, similar to that of our genetic deletion alone (Figure 4). This indicates that maternal separation as an environmental stressor alone has a similar impact on myelination compared to the genetic deletion alone, at least during adolescence. Additionally, our two-hit model allowed us to look at the impact of both maternal separation and genetic deletion on myelination. We observed similar reductions in myelination as the genetic deletion and environmental separation alone (Figure 4). All three conditions showed decreased myelination compared to non-separated controls.

### Effects on earlier developmental stages

We also wanted to examine if there are visible differences in the OL lineage and myelination during the prodromal phase of schizophrenia, examining mice at P14. We found a significant decrease in proliferation, as measured by the density of EdU+Olig2+ OPCs, only in the mPFC (Figure 5). This suggests that this brain region is especially vulnerable. We see no differences in immature or mature oligodendrocyte density as measured by BCAS+/Olig2+ (Figure 6) or CC1+/Olig2+ OLs (Figure 7). Additionally, we found little differences in myelination in P14 mice, except a significant reduction of MBP area fraction in the dlPFC (Figure 8). We do see reduced myelin process lengths in the mPFC, again demonstrating this regions as particularly vulnerable (Figure 8).

**Figure 5.**
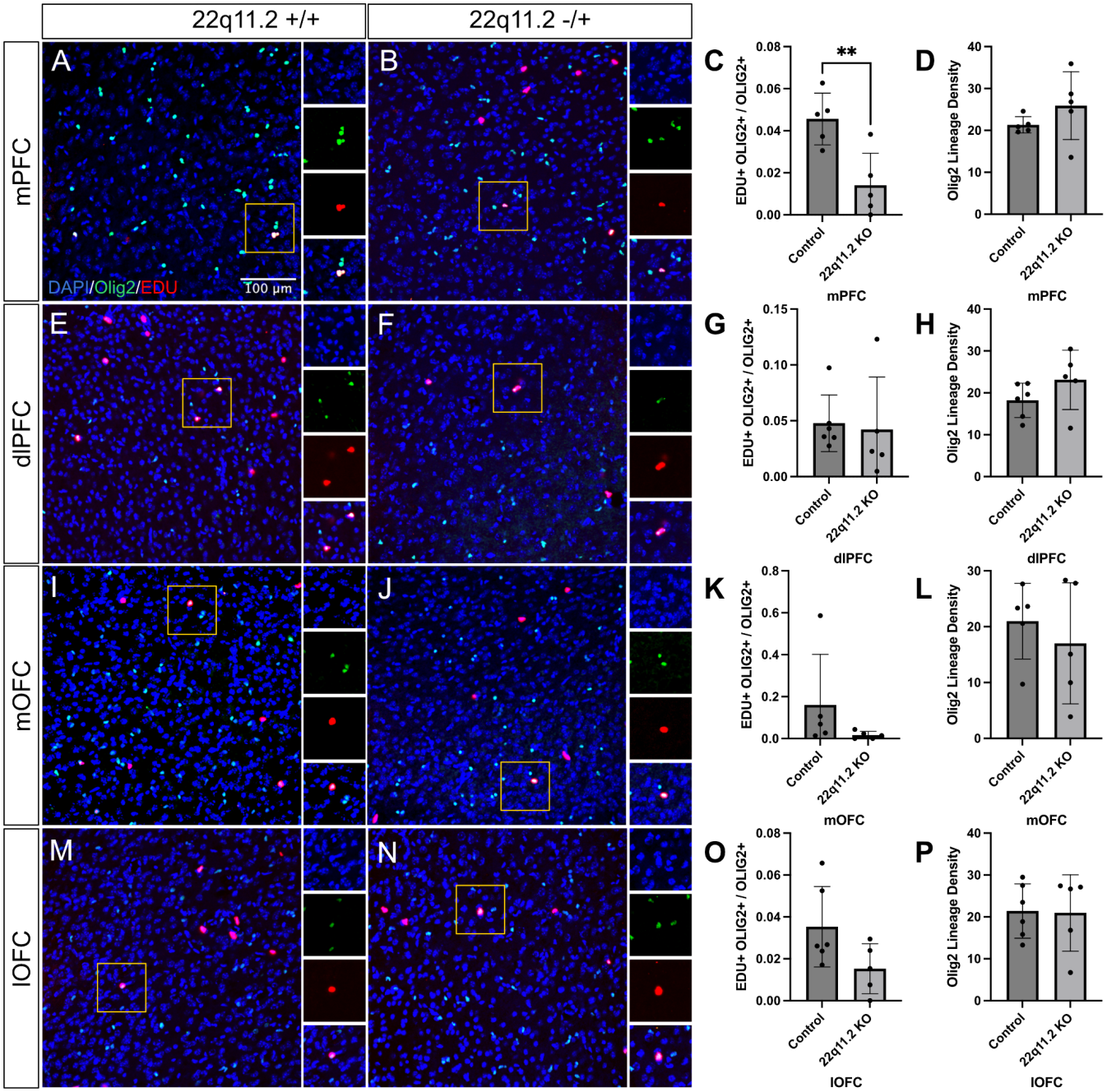
Early in Postnatal Development, 22q11.2 Deletion Mice Have a Decrease in OPCs Only in the Medial Prefrontal Cortex. Representative coronal sections showing DAPI (blue), Olig2 (green), and click-it chemistry EdU (red) staining in P14 WT (n=5/6) and 22q11.2 KO (n=5) mice across the medial prefrontal cortex (A, B), dorsolateral prefrontal cortex (E, F), medial orbitofrontal cortex (I, J), and lateral orbitofrontal cortex (M, N). (C, G, K, O) Ratio of EdU+Olig2+ cells to total Olig2+ cells. (D, H, L, P) Olig2+ lineage density. Data presented as mean ± SEM, unpaired two-tailed t-test: (C) t=3.597, df=8, p=0.0070; (D) t=1.231, df=8, p=0.2534; (G) Kolmogorov-Smirnov test d=0.6000, p=0.2100; (H) t=1.437, df=9, p=0.1844; (K) Kolmogorov-Smirnov test d=0.6000, p=0.2100; (L) t=0.6930, df=8, p=0.5079; (O) t=2.021, df=9, p=0.039; (P) t=0.09871, df=9, p=0.9235.

**Figure 6.**
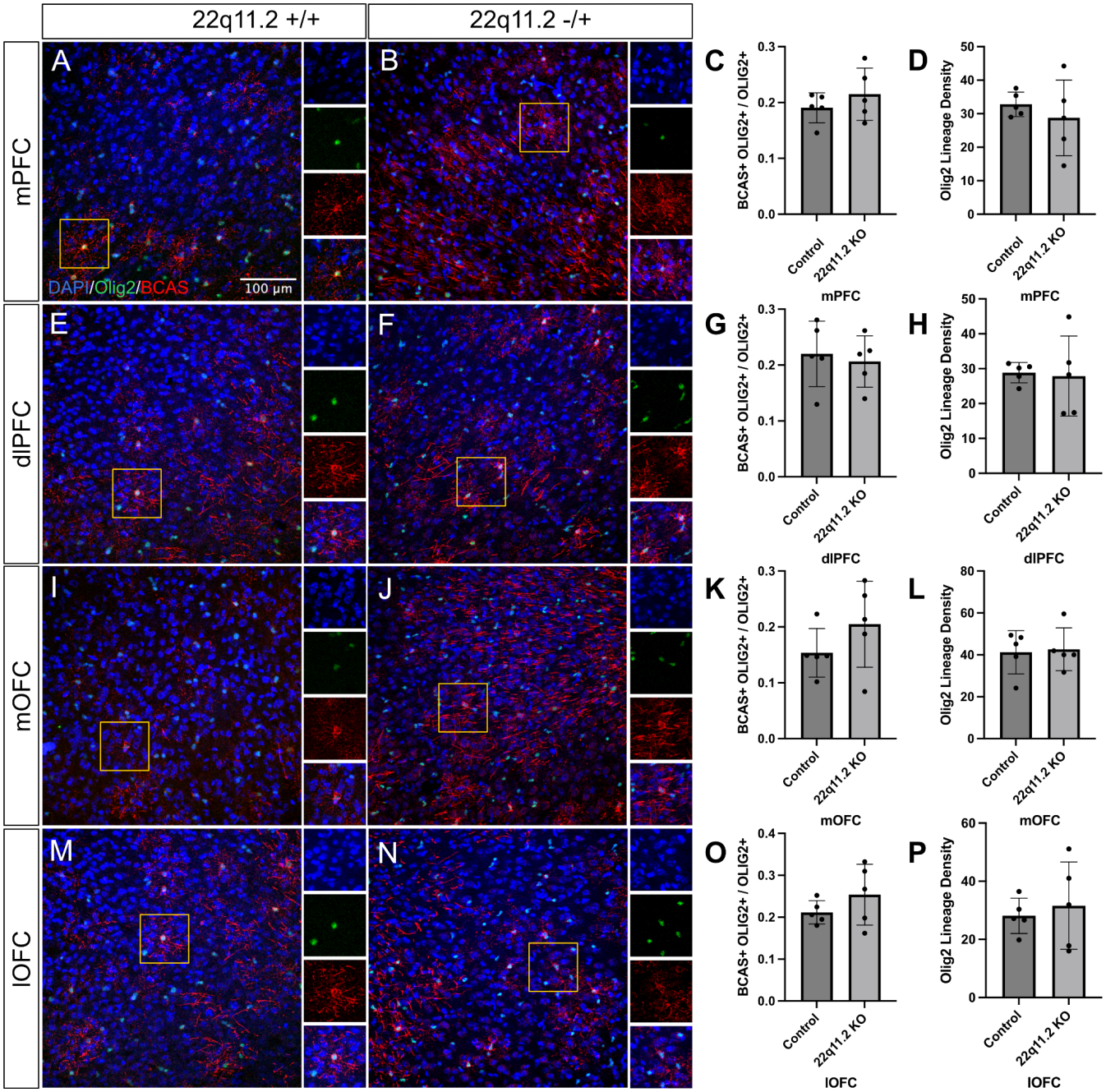
Early in Postnatal Development, 22q11.2 Deletion Mice Have No Change in Immature Oligodendrocytes in the Prefrontal Cortex. Representative coronal sections showing DAPI (blue), Olig2 (green), and BCAS (red) staining in P14 WT (n=5) and 22q11.2 KO (n=5) mice across the medial prefrontal cortex (A, B), dorsolateral prefrontal cortex (E, F), medial orbitofrontal cortex (I, J), and lateral orbitofrontal cortex (M, N). (C, G, K, O) Ratio of BCAS+Olig2+ cells to total Olig2+ cells. (D, H, L, P) Olig2+ lineage density. Data presented as mean ± SEM, unpaired two-tailed t-test: (C) t=1.005, df=8, p=0.3441; (D) t=0.7633, df=8, p=0.4672; (G) t=0.4103, df=8, p=0.6923; (H) t=0.1845, df=8, p=0.8582; (K) t=1.295, df=8, p=0.2313; (L) t=0.2179, df=8, p=0.8329; (O) t=1.217, df=8, p=0.2584; (P) t=0.4818, df=8, p=0.6429.

**Figure 7.**
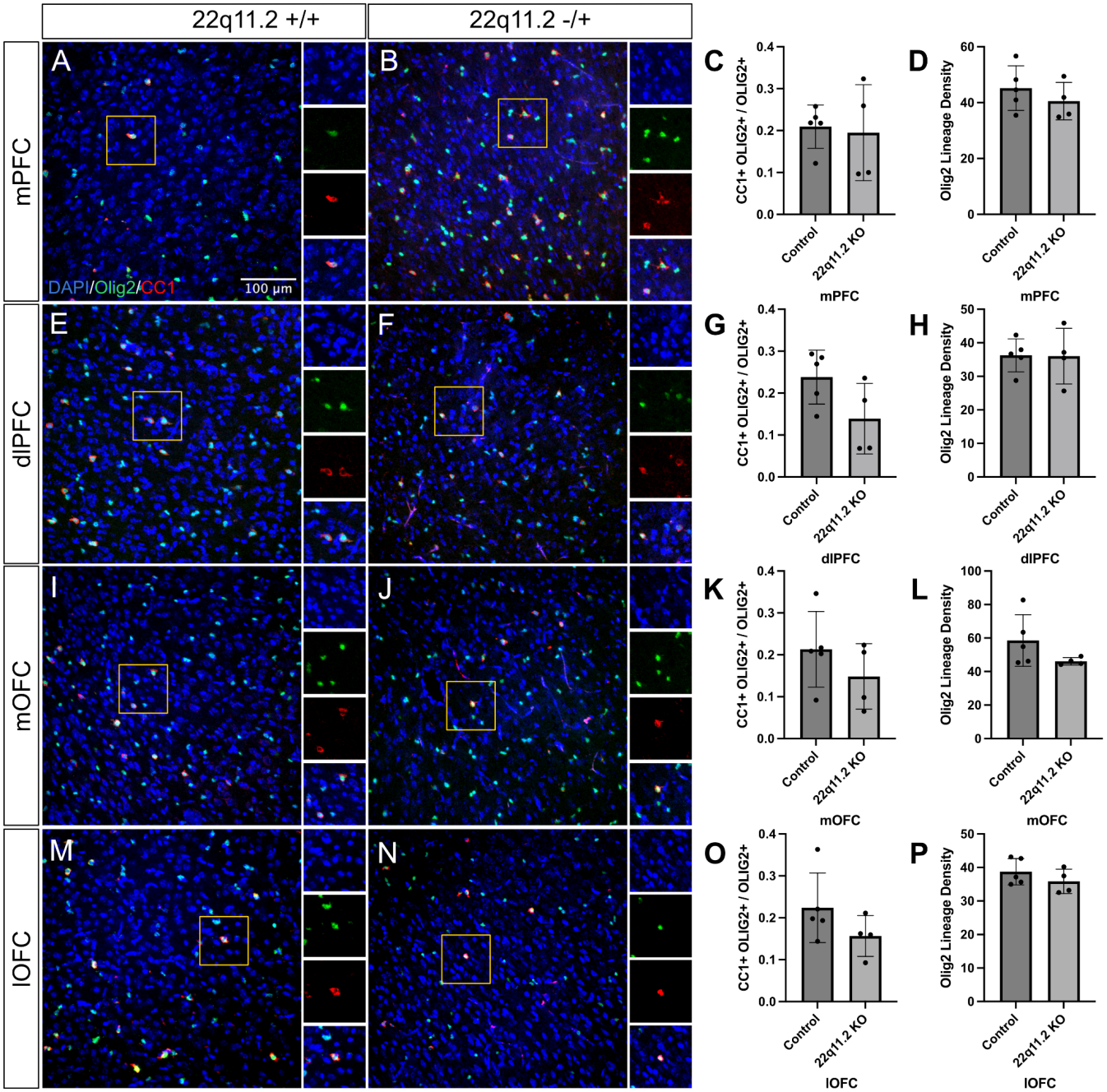
Early in Postnatal Development, 22q11.2 Deletion Have No Change in Mature Oligodendrocytes in the Prefrontal Cortex. Representative coronal sections showing DAPI (blue), Olig2 (green), and CC1 (red) staining in P14 WT (n=5) and 22q11.2 KO (n=4) mice across the medial prefrontal cortex (A, B), dorsolateral prefrontal cortex (E, F), medial orbitofrontal cortex (I, J), and lateral orbitofrontal cortex (M, N). (C, G, K, O) Ratio of CC1+Olig2+ cells to total Olig2+ cells. (D, H, L, P) Olig2+ lineage density. Data presented as mean ± SEM, unpaired two-tailed t-test: (C) t=0.2586, df=7, p=0.8034; (D) t=0.9347, df=7, p=0.3811; (G) t=2.015, df=7, p=0.0838; (H) t=0.05189, df=7, p=0.9601; (K) t=1.135, df=7, p=0.2939; (L) t=1.577, df=7, p=0.1587; (O) t=1.430, df=7, p=0.1957; (P) t=1.128, df=7, p=0.2963.

**Figure 8.**
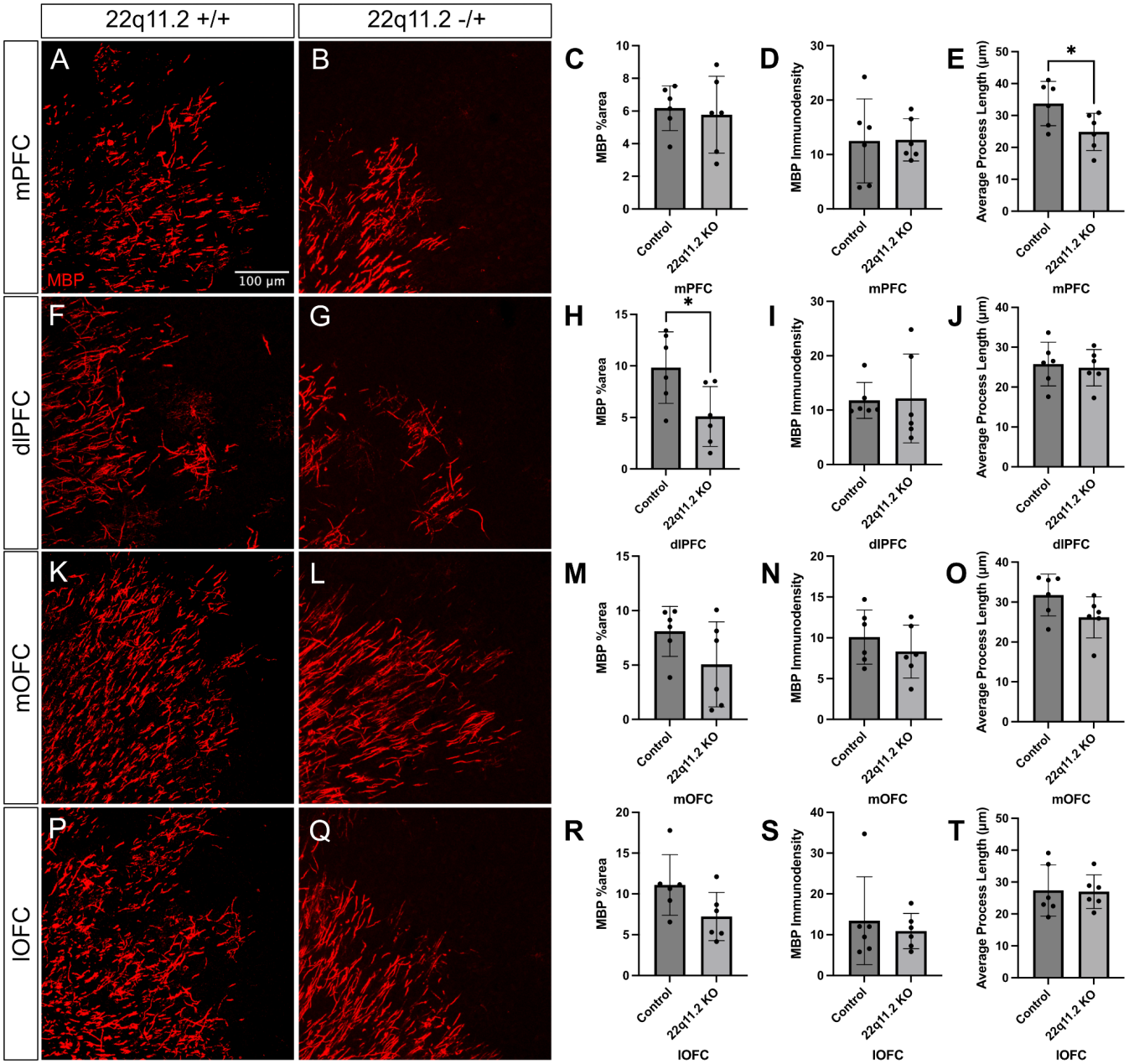
Early in Postnatal Development, 22q11.2 Deletion Mice Have Myelination Defects in the Prefrontal Cortex. Representative coronal sections showing MBP (red) staining in P14 WT (n=6) and 22q11.2 KO (n=6) mice across the medial prefrontal cortex (A, B), dorsolateral prefrontal cortex (F, G), medial orbitofrontal cortex (K, L), and lateral orbitofrontal cortex (P, Q). (C, H, M, R) MBP area fraction. (D, I, N, S) MBP immunodensity. (E, J, O, T) MBP average process length (μm). Data presented as mean ± SEM, unpaired two-tailed t-test: (C) t=0.3625, df=10, p=0.7245; (D) t=0.05307, df=10, p=0.9587; (E) t=3.664, df=10, p=0.0370; (H) t=2.570, df=10, p=0.0279; (I) Kolmogorov-Smirnov test d=0.6667, p=0.1429; (J) t=0.3185, df=10, p=0.7567; (M) t=1.636, df=10, p=0.1329; (N) t=0.9370, df=10, p=0.3708; (O) t=0.2572, df=10, p=0.0924; (R) t=1.994, df=10, p=0.0741; (S) Kolmogorov-Smirnov test d=0.1667, p=>0.9999; (T) t=0.1017, df=10, p=0.9210.

## Discussion

Our results demonstrate that the DiGeorge 22q11.2 deletion model has significantly altered oligodendrocyte (OL) maturation and myelination in the prefrontal cortex (PFC). We found that in the prodromal phase (P14) there are slight differences in oligodendrocyte precursor cells (OPCs) and myelin structure, specifically in the medial PFC, demonstrating early differences caused by genetic alterations in a vulnerable region of the PFC. In late adolescence (P28), we found severe differences in oligodendrocyte maturation, including increased immature and mature cell density. Along with these abnormalities in maturation, we found a drastic decrease in myelination in all regions of the prefrontal cortex. In addition, we determined the impact of an additional environmental stressor of maternal separation to examine a two-hit model similar to that hypothesized for humans with schizophrenia. We found a similar decrease in myelination compared to the genetic-only model in both the stressor-only model and two-hit model. This suggests that both genetic deficits and environmental stress can lead to similar defects in the development of OLs and their ability to successfully myelinate axons in the PFC.

Our work highlights the impact of the 22q11.2 deletion on oligodendrocyte lineage and function. Previous research demonstrated abnormalities in the morphology of post-mortem OPCs in the hippocampus, cortex, and amygdala^18^. Another study demonstrated a decrease in the density of OLs but not OPCs in post-mortem tissue^19^. These studies suggest abnormalities in maturation of OLs. Our results align with past work, demonstrating decreased myelination in the PFC. We discovered increased density in mature and immature OLs, a new progression highlighting abnormalities of oligodendrocyte maturation. These results could be specific to the 22q11.2 deletion and should be further explored in comparison to other mouse models of genetic risk that phenocopy schizophrenia. We found no change in proliferating OPCs and increased density of mature OLs, which is different from previous research mentioned above. Past results in this model have also had inconsistent findings regarding overall OL density changes, and our results suggest that there are no alterations in the overall density.

It is possible that the increase in density is due to rapid maturation of OLs, which disrupts signaling pathways and prevents OLs from properly myelinating. Past research has shown increased junctions and branching of OLs^20^, which provides evidence towards rapid maturation. It is also possible that there are factors directly impacting the ability of OLs to myelinate, such as faulty wrapping of axons. This would need to be addressed in future research. Our results demonstrate that there are specific abnormalities within OLs, providing an explanation for the decreased or altered connectivity within the prefrontal cortex and its pathways to the rest of the brain found in humans with schizophrenia^21^. This brain circuit is especially relevant to the cognitive symptoms of schizophrenia, which may explain symptoms in patients with schizophrenia related to deficits in cognition, attention, and memory.

The lifetime prevalence of schizophrenia is around 1%, and its symptoms heavily impact one’s daily life and relationships. Evidence for genetic vulnerability for schizophrenia is fairly strong, with around 70% heritability. The 22q11 deletion syndrome is present in around 1% of individuals with schizophrenia, presenting a strong model to understand the genetic impacts of the disease. Treatments are heavily focused on resolving positive symptoms of schizophrenia, but leave the cognitive symptoms largely untreated. A better understanding of the factors that play a role in the cognitive deficits could provide future avenues for treatment to improve the lives of many individuals with schizophrenia. In addition, a stronger understanding of the role of OL changes in psychiatric diseases provides opportunities for novel treatments in the future.

## Footnotes

1 [Schizophrenia and 22q11.2 Deletion Syndrome - PMC](https://pmc.ncbi.nlm.nih.gov/articles/PMC3129332/)

2 [Schizophrenia and 22q11.2 Deletion Syndrome - PMC](https://pmc.ncbi.nlm.nih.gov/articles/PMC3129332/)

3 [In vivo evidence of microstructural hypo-connectivity of brain white matter in 22q11.2 deletion syndrome | Molecular Psychiatry](https://www.nature.com/articles/s41380-023-02178-w)

4 [A population study of chromosome 22q11 deletions in infancy - PMC](https://pmc.ncbi.nlm.nih.gov/articles/PMC1717723/)

5 [Schizophrenia and 22q11.2 Deletion Syndrome - PMC](https://pmc.ncbi.nlm.nih.gov/articles/PMC3129332/)

6 [Tbx1 Heterozygosity in the Oligodendrocyte Lineage Shifts Myelinated Axon Composition in the Mouse Fimbria Without Behavioral Impairments - PMC](https://pmc.ncbi.nlm.nih.gov/articles/PMC1717723/)

7 [Developmentally dynamic, non-convergent transcriptomic profiles in CNV models for schizophrenia risk | Molecular Psychiatry](https://www.nature.com/articles/s41380-026-03679-0#Sec2)

8 [Risk factors and the evolution of psychosis in 22q11.2 deletion syndrome: a longitudinal 2-site study - PubMed](https://pubmed.ncbi.nlm.nih.gov/24157393/)

9 https://cshperspectives.cshlp.org/content/16/6/a041358

10 [Adolescence is associated with genomically patterned consolidation of the hubs of the human brain connectome - PMC](https://pmc.ncbi.nlm.nih.gov/articles/PMC4987797/)

11 [Prolonged myelination in human neocortical evolution - PMC](https://pmc.ncbi.nlm.nih.gov/articles/PMC3478650/)

12 [The mechanistic target of rapamycin pathway downregulates bone morphogenetic protein signaling to promote oligodendrocyte differentiation - PubMed](https://pubmed.ncbi.nlm.nih.gov/31904150/)

13 [Tbx1, a gene encoded in 22q11.2 copy number variant, is a link between alterations in fimbria myelination and cognitive speed in mice | Molecular Psychiatry](https://www.nature.com/articles/s41380-021-01318-4)

14 [Tbx1, a gene encoded in 22q11.2 copy number variant, is a link between alterations in fimbria myelination and cognitive speed in mice | Molecular Psychiatry](https://www.nature.com/articles/s41380-021-01318-4)

15 [The dual hit hypothesis of schizophrenia: Evidence from animal models - ScienceDirect](https://www.sciencedirect.com/science/article/pii/S014976342100467X?via%3Dihub)

16 [Neonatal Maternal Separation Impairs Prefrontal Cortical Myelination and Cognitive Functions in Rats Through Activation of Wnt Signaling | Cerebral Cortex | Oxford Academic](https://academic.oup.com/cercor/article-abstract/27/5/2871/3056363)

17 [Early-life stress impairs postnatal oligodendrogenesis and adult emotional behaviour through activity-dependent mechanisms | Molecular Psychiatry](https://www.nature.com/articles/s41380-019-0493-2)

18 [Pathological oligodendrocyte precursor cells revealed in human schizophrenic brains and trigger schizophrenia-like behaviors and synaptic defects in genetic animal model](https://www.nature.com/articles/s41380-022-01777-3)

19 [Differentiation of Oligodendrocyte Precursors is Impaired in the Prefrontal Cortex in Schizophrenia](https://pmc.ncbi.nlm.nih.gov/articles/PMC4681621/#S2)

20 [iPSC-modelling reveals genetic associations and morphological alterations of oligodendrocytes in schizophrenia](https://www.nature.com/articles/s41398-025-03509-x#Sec20)

21 [In vivo evidence of microstructural hypo-connectivity of brain white matter in 22q11.2 deletion syndrome | Molecular Psychiatry](https://www.nature.com/articles/s41380-023-02178-w)

